# Evolutionary history of Carnivora (Mammalia, Laurasiatheria) inferred from mitochondrial genomes

**DOI:** 10.1101/2020.10.05.326090

**Authors:** Alexandre Hassanin, Géraldine Véron, Anne Ropiquet, Bettine Jansen van Vuuren, Alexis Lécu, Steven M. Goodman, Jibran Haider, Trung Thanh Nguyen

## Abstract

The order Carnivora, which currently includes 296 species classified into 16 families, is distributed across all continents. The phylogeny and the timing of diversifications are still a matter of debate.

Here, complete mitochondrial genomes were analysed to reconstruct the phylogenetic relationships and to estimate divergence times among species of Carnivora. We assembled 51 new mitogenomes from 13 families, and aligned them with available mitogenomes by selecting only those showing more than 1% of nucleotide divergence and excluding those suspected to be of low-quality or from misidentified taxa. Our final alignment included 220 taxa representing 2,442 mitogenomes. Our analyses led to a robust resolution of suprafamilial and intrafamilial relationships. We identified 22 fossil calibration points to estimate a molecular timescale for carnivorans. According to our divergence time estimates, crown carnivorans appeared during or just after the Early Eocene Climatic Optimum; all major groups of Caniformia (Cynoidea/Arctoidea; Ursidae; Musteloidea/Pinnipedia) diverged from each other during the Eocene, while all major groups of Feliformia (Nandiniidae; Feloidea; Viverroidea) diversified more recently during the Oligocene, with a basal divergence of *Nandinia* at the Eocene/Oligocene transition; intrafamilial divergences occurred during the Miocene, except for the Procyonidae, as *Potos* separated from other genera during the Oligocene.

## Introduction

The order Carnivora is composed of 296 extant species (IUCN, 2020) currently ranged into two suborders: the Caniformia which includes nine families, namely the Canidae (dog-like species), Ailuridae (red panda), Mephitidae (skunks and stink badgers), Mustelidae (weasels, badgers, martens, otters, etc.), Odobenidae (walrus), Otariidae (eared seals), Phocidae (earless seal), Procyonidae (raccoons, coatis, kinkajous, etc.), and Ursidae (bears); and the Feliformia which is represented by seven families, namely the Felidae (cat-like species), Eupleridae (Malagasy carnivorans), Herpestidae (mongooses), Hyaenidae (hyenas), Nandiniidae (African palm civet), Prionodontidae (Asiatic linsangs), and Viverridae (civets and genets). The oldest known fossils of Carnivora have been found in the late Paleocene; they belong to the extinct families Miacidae and Viverravidae, and have small body-size, comparable to extant weasels and martens (Spaulding & Flynn, 2012; Solé et al., 2014; 2016). The timing of the emergence of the crown carnivorans and their relationships to Paleocene and Eocene fossils are still unresolved. However, there is consensus for supporting two higher fossil taxa: the Carnivoraformes, which are composed of the crown group plus the stem family Miacidae, which is probably paraphyletic; and the Carnivoramorpha, which groups the Carnivoraformes and the Viverravidae (Flynn et al., 2010; Solé et al., 2016).

The phylogeny of several carnivoran families has been extensively studied based on mitochondrial and nuclear data, including the Felidae, the Mustelidae, and the Ursidae (Kutschera et al., 2014; Li et al., 2016; Law et al., 2018), while other families remain poorly studied, specifically the Eupleridae, Herpestidae, Mephitidae, Procyonidae, and Viverridae. The phylogeny and timescale of diversification of the Carnivora have been studied by Eizirik et al. (2010) using a molecular supermatrix of 7,765 base pairs (bp) containing 14 nuclear genes for 50 species, which represents less than 17% of the species diversity of the order Carnivora. The evolutionary history of carnivorans has also been inferred using a supertree approach by Nyakatura & Bininda-Emonds (2012). These two studies were based on different methods and data, and different fossils were used as calibration points. Although several nodes showed similar ages in the two studies, such as the most recent common ancestor (MRCA) of the Carnivora (59.2 versus 64.9 Mya), Arctoidea (42.6 vs 47.8 Mya) and Pinnipedia (24.5 vs 22.4 Mya), some nodes were highly discordant, including the MRCA of the Caniformia (48.2 vs 61.2 Mya), Feliformia (44.5 vs 53.2 Mya), Feloidea (Felidae + Prionodontidae) (33.3 vs 52.9 Mya) and Canidae (7.8 vs 16.3 Mya). Most other studies to date have focused on either the Caniformia (e.g. Law et al., 2018) or the Feliformia (e.g. Zhou et al., 2017) and the timing of diversification in some families appears highly elusive or uncertain in the absence of a molecular timescale based on a high diversity of species from all carnivoran families.

With the development of next-generation sequencing (NGS) technologies, the number of mitochondrial genomes available in the international nucleotide databases has considerably increased during the last decades (Coissac et al., 2016). For the order Carnivora, there are currently more than 2,400 complete mitogenomes, and some of these were sequenced from Pleistocene fossils, such as polar bear (Lindqvist et al., 2010), giant short-faced bears (Mitchell et al., 2016), cave lion (Barnett et al., 2016) or saber-toothed cats (Paijmans et al., 2017). This huge and diversified dataset offers an excellent opportunity to better understand the evolutionary history of the order Carnivora, as problematic sequences and taxonomic issues can be more easily detected, and more importantly, as many fossils can be included as calibration points for estimating divergence times. The latter aspect is particularly relevant considering the paleontological record of Carnivora has been significantly improved over the last ten years, with the discovery of several key fossils (e.g. Abella et al., 2012; Grohé et al., 2013; Tseng et al., 2014; Berta et al., 2018; Valenciano et al., 2020).

Here we analysed complete mitochondrial genomes to reconstruct phylogenetic relationships among carnivorans and to estimate divergence times. We sequenced 43 mitogenomes using various methods (Sanger sequencing of PCR products, or next-generation sequencing of long PCR products or Illumina shotgun sequencing). We also assembled eight mitogenomes from Sequence Read Archive (SRA) data. The 51 new mitogenomes belong to several families, namely Canidae (2), Eupleridae (6), Felidae (4), Herpestidae (12), Hyaenidae (1), Mustelidae (10), Otariidae (2), Phocidae (1), Prionodontidae (1), Procyonidae (3), Ursidae (1), Viverridae (7) for Carnivora, as well as Tapiridae (1) for the order Perissodactyla, which was included as an outgroup. These new mitogenomes were compared to all mitogenomes available for Carnivora in public repositories. At the intraspecific level, we selected only mitogenomes that were separated by more than 1% of nucleotide divergence and excluded those suspected to be of low-quality or from misidentified taxa. Our final alignment includes 220 taxa, which represent 2,442 mitogenomes. Our main objective was to build one of the largest time-trees of Carnivora estimated using a selection of fossil calibration points in order to provide further insights into the evolution of this broadly distributed and morphologically diverse taxonomic group.

## Material and Methods

### DNA extraction, amplification, sequencing and mitogenome assembly

Total DNA was extracted from cells, muscle, or skin samples using the DNeasy Blood and Tissue Kit (Qiagen, Hilden, Germany). Details on the 43 samples extracted for this study are given in S1 Appendix. The mitochondrial genomes were sequenced using one of the three following approaches: Sanger sequencing of ≈ 20 overlapping PCR products (length between 700 and 2000 bp); next-generation sequencing (NGS) of five overlapping long PCR products of around 4-5 kb; and shotgun Illumina sequencing.

In the first approach, PCR amplifications were carried out as described in Hassanin et al. (2012) using the primers listed in S2 Appendix. The amplicons were then sequenced in both directions by Eurofins MWG Operon (Ebersberg, Germany). Genomes were assembled with electropherograms of overlapping amplicons using Sequencher 5.1 (Gene Codes Corporation, Ann Arbor, MI, USA).

In the second approach, five overlapping PCR products of around 4-5 kb were amplified as described in Hassanin et al. (2020), and they were sequenced at the “Service de Systématique Moléculaire” (UMS CNRS 2700, MNHN, Paris, France) using the Ion Torrent Personal Genome Machine (Thermo Fisher Scientific, Waltham, MA, USA).

The third approach was based on shotgun Illumina sequencing. DNA samples were quantified with a Qubit® 2.0 Fluorometer using the Qubit dsDNA HS Assay Kit (Thermo Fisher Scientific, Waltham, MA, USA). Libraries were prepared using the TruSeq® Nano DNA Library Prep kit (Illumina, San Diego, CA, USA) after pooling 150 ng of total DNA of 10 species belonging to distant taxonomic groups (i.e. different phyla, classes, orders or families). Libraries were sequenced at the “Institut du Cerveau et de la Moelle épinière” (Paris, France) using a NextSeq® 500 system and the NextSeq 500 High Output Kit v2 (300 cycles) (Illumina).

The NGS reads generated with either Ion Torrent or Illumina sequencers were assembled by baiting and iterative mapping approach on Geneious® 10.2.2 (Biomatters Ltd., Auckland, New Zealand) using available mitochondrial references, including cytochrome b, cytochrome c oxidase subunit I, 12S and 16S rRNA genes. The 43 new mitochondrial genomes generated for this study were annotated on Geneious and deposited in GenBank under accession numbers XXXXXX-XXXXXX.

Eight mitochondrial genomes were assembled from SRA downloaded from NCBI for the following species: *Bassaricyon neblina* (SRX1097850), *Bassariscus sumichrasti* (SRX1099089), *Helogale parvula* (SRR7637809), *Lontra canadensis* (SRR10409165), *Mirounga angustirostris* (SRR10331586), *Mungos mungo* (SRR7704821), *Prionodon linsang* (ERR2391707), and *Zalophus wollebaeki* (SRR4431565).

### Mitochondrial alignments

The 51 mitochondrial genomes assembled in this study were compared with other genomes available in the NCBI nucleotide database (see accession numbers in S1 Appendix) for the different families. At the intraspecific level, we selected only mitochondrial haplotypes separated by more than 1% of divergence. Our final alignment includes 218 mitogenomes of Carnivora representing a large taxonomic diversity in the 16 following families (in parentheses number of mitgenomes): Ailuridae (2), Canidae (21), Eupleridae (7), Felidae (46), Herpestidae (15), Hyaenidae (4), Mephitidae (3), Mustelidae (39), Nandiniidae (1), Odobenidae (1), Otariidae (12), Phocidae (30), Prionodontidae (2), Procyonidae (6), Ursidae (21), and Viverridae (19). A tapir (*Tapirus terrestris*) and a pangolin (*Phataginus tricuspis*) were used to root the carnivoran tree because they represent two Laurasiatherian orders, Perissodactyla and Pholidota, respectively, which were shown to be closely related to Carnivora in previous molecular studies (e.g. Murphy et al., 2001; Meredith et al., 2011).

The 220 mitochondrial genomes were aligned using AliView 1.22 (Larsson, 2014). Ambiguous regions for primary homology were excluded from the alignment. To limit the impact of missing data, we also removed from the alignment all indels (insertions or deletions) detected in only one genome. The final alignment, named *mtDNA*, contains 220 taxa and 14,892 nucleotide sites. Two other datasets were used for the analyses: (1) the *mtDNA-Tv* dataset (transversions only), which corresponds to the *mtDNA* dataset in which the nucleotide G was replaced by A and the nucleotide T by C; and (2) the *PCG-DNA* dataset (10,809 nt) in which all regions other than protein-coding genes were removed (i.e. control region, 12S and 16S rRNA genes and tRNA genes), as well as the *ND6* gene (because it is located on the opposite strand of other protein-coding genes). The three datasets used in this study are available at https://osf.io/XXX.

### Analysis of base composition

The alignment of the protein-coding genes of 220 mitochondrial genomes (*PCG-DNA* dataset) was used to calculate the frequency of the four bases (A, C, G and T) at each of the three codon-positions (S3 Appendix). The twelve variables measured were then summarized by a principal component analysis (PCA) using the FactoMineR package (Lê et al., 2008) in R version 3.5.3 (from http://www.R-project.org/). The strand bias in nucleotide composition was studied at third codon-positions of the *PCG-DNA* dataset by calculating the relative frequencies of A and T nucleotides (AT3 skew = [A - T] / [A + T]) and the relative frequencies of C and G nucleotides (CG3 skew = [C - G] / [C + G]) (Lobry, 1995; Hassanin et al., 2005; Arabi et al., 2010).

### Phylogenetic analyses

Two datasets (*mtDNA* and *mtDNA-Tv*) were analysed with probabilistic methods for tree reconstruction using the resources available from the CIPRES Science Gateway (Miller et al., 2010). The Bayesian analyses were done with MrBayes 3.2.7 (Ronquist et al., 2012) and the two following models: GTR+I+G for the *mtDNA* dataset, and JC69+I+G for the *mtDNA-Tv* dataset. The posterior probabilities (PP) were calculated using 10,000,000 Metropolis-coupled MCMC generations, tree sampling every 1000 generations, and a burn-in of 25%.

To examine the phylogenetic signal along the *mtDNA* dataset, we also performed Bayesian analyses (with the same parameters) on 10 half-overlapping sub-datasets (i–x) of the about the same length (i.e., 2978 or 2980 nt), corresponding to the following positions: (i) 1-2978; (ii) 1489-4466; (iii) 2979-5956; (iv) 4467-7444; (v) 5957-8934; (vi) 7445-10422; (vii) 8935-11912; (viii) 10423-13400; (ix) 11913-14892; and (x) 13401-14892 + 1-1488. The use of half-overlapping sub-datasets (sliding window of ≈ 2978 nt) implies that all nucleotide sites of the total *mtDNA* alignment are represented twice in these Bayesian analyses. The lists of bipartitions obtained from Bayesian analyses of the 10 sub-datasets were transformed into a weighted binary matrix for supertree construction using SuperTRI v57 (Ropiquet et al., 2009). Each binary character corresponds to a node, which was weighted according to its frequency of occurrence in one of the 10 lists of bipartitions. In this manner, the SuperTRI method takes into account both principal and secondary signals, because all phylogenetic hypotheses found during the 10 Bayesian analyses were used for the calculation of the following two reliability indices for each node of interest: (1) the supertree bootstrap percentages (SBP) were obtained from PAUP 4* version 4b10 (Swofford, 2003) after 1000 bootstrap replicates of the MRP (Matrix Representation with Parsimony) matrix of 3,398 binary characters (reconstructed under SuperTRI v57); and (2) the mean posterior probabilities percentages (MPP) were directly calculated on SuperTRI v57. The SBP and MPP values were reported on the Bayesian tree found with the total alignment of 14,892 nt. Here, the SuperTRI analyses were conducted to test for phylogenetic signal along the mtDNA genome. If a robust node in the Bayesian tree (PP ≥ 0.95) is recovered with high SBP (≥ 95%) and MPP values (≥ 0.80), it means that the signal is present all along the mtDNA genome. If a node in the Bayesian tree is recovered with low MPP values (< 0.50), it means that the signal is weak or confined to a few fragments of the mtDNA genome. If there is a robust topological conflict between Bayesian and SuperTRI results, it suggests that at least one of the studied genomes was partially contaminated by a mitochondrial DNA sequence from another species or by a nuclear DNA sequence of mitochondrial origin (Numt). An example was published in Hassanin et al. (2010) for the mitochondrial genomes of domestic goat.

### Molecular dating

Divergence times were estimated on the CIPRES Science Gateway (Miller et al., 2010) using the *mtDNA* dataset and the Bayesian approach implemented in BEAST v.2.4.7 (Bouckaert et al., 2014). Twenty-two calibration points were selected for molecular dating (**Table 1**). Most of these were interpreted from the fossil record using maximum (Max) and minimum (Min) ages. We applied two strategies for fossil calibration: (1) a uniform distribution between Max and Min on the calibrated node ages; or (2) a log-normal distribution on the calibrated node ages using Min as offset, M = Max - Min / 4, and S = 0.926 (to match the 97.5% quantile to Max) (see details in Table 1). The second strategy relies on the fact that minimum ages are generally more accurate and reliable than maximum ages because younger fossils are more abundant and more accurately dated than older fossils as a consequence of taphonomic processes and dating methods (Crees et al., 2019). We applied a GTR + I + G model for the *mtDNA* alignment (with a proportion of invariants of 0.425) and a relaxed-clock model with uncorrelated lognormal distribution for substitution rates. Node ages were estimated using a calibrated Yule speciation prior and 2.10^8^ generations, with tree sampling every 2,000 generations, and a burn-in of 25% generations. MCMC mixing efficiency and convergence were assessed using the ESS values (>200) in Tracer v.1.7.1. The chronogram was reconstructed with TreeAnnotator, which is included in the BEAST package (Bouckaert et al., 2014).

**Table 1.** Maximum (Max) and minimum (Min) age calibrations (in millions of years ago, Mya) used for molecular dating analyses (with either uniform or log-normal prior distributions [U/L] on calibrated node ages interpreted from the fossil record).

| Most recent common ancestor | Max | Distribution | Min | References and justification |
| --- | --- | --- | --- | --- |
| **Ferae** | 85 | N, M = 75 | 65 | www.timetree.org ; split *Canis* / *Manis* (13 studies) |
| **Caniformia** |  |  |  |  |
| **Canidae** |  |  |  |  |
| (*Canis*, *Cuon*, *Lycaon*) | 12 | U/L, M = 1.5 | 6 | Tedford et al. (2009) |
| (*Vulpes*, *Nyctereutes*) | 34 | U/L, M = 6.25 | 9 | Tedford et al. (2009) |
| **Ursidae** | 34 | U/L, M = 5.75 | 11 | Abella et al. (2012), Banyue & Zhanxiang (2003), Bonis et al. (2019) |
| (*Arctotherium*, *Tremarctos*) | 10 | U/L, M = 5.75 | 3 | Soibelzon et al. (2005) |
| *Ursus maritimus*† from Svalbard (GU573488) | 0.11 | N, M = 0.12 | 0.13 | Lindqvist et al. (2010) |
| **Pinnipedia** | 34 | U/L, M = 3.75 | 19 | Rybczynski et al. (2009), Berta et al. (2018), Dewaele et al. (2018) |
| **Otarioidea** | 23 | U/L, M = 2.00 | 15 | Berta et al. (2018) |
| **Phocidae** | 23 | U/L, M = 2.75 | 12 | Berta et al. (2018) |
| **Mustelidae** |  |  |  |  |
| (Guloninae ... Mustelinae) | 27.6 | U/L, M = 2.825 | 16.3 | Sato et al. (2009), Valenciano et al. (2020) |
| (Lutrinae, Ictonychinae) | 27.6 | U/L, M = 3.85 | 12.2 | Sato et al. (2009), Grohé et al. (2013) |
| **Feliformia** |  |  |  |  |
| **Feloidea** | 34 | U/L, M = 3.5 | 20 | Werdelin et al. (2010) |
| **Felidae** | 20 | U/L, M = 1.5 | 14 | Werdelin et al. (2010) |
| (*Acinonyx*, *Puma*) | 14 | U/L, M = 2.65 | 3.4 | Werdelin et al. (2010), Werdelin & Peigné (2010) |
| (*Leptailurus* ... *Profelis*) | 14 | U/L, M = 2.5 | 4 | Werdelin & Peigné (2010) |
| (*Caracal*, *Profelis*) | 14 | U/L, M = 2.5 | 4 | Werdelin et al. (2010) |
| (*Panthera*, *Neofelis*) | 14 | U/L, M = 2.0125 | 5.95 | Tseng et al. (2014) |
| **Viverroidea** |  |  |  |  |
| **Viverridae** |  |  |  |  |
| (Genettinae, Viverrinae) | 34 | U/L, M = 5.00 | 14 | Werdelin & Peigné (2010) |
| **(Herpestidae, Eupleridae)** | 34 | U/L, M = 3.375 | 20.5 | Werdelin & Peigné (2010) |
| (*Helogale*, *Crossarchus*) | 20.5 | U/L, M = 3.625 | 6 | Werdelin & Peigné (2010) |
| (*Galerella* ... *Cynictis*) | 20.5 | U/L, M = 3.275 | 7.4 | Werdelin & Peigné (2010) |
| **Hyaenidae** |  |  |  |  |
| (*Hyaena*, *Parahyaena*) | 9.5 | U/L, M = 1.475 | 3.6 | Ferretti (2007), Werdelin & Peigné (2010) |

Abbreviations: L = Log Normal; M = Mean age; N = Normal; U: Uniform.

## Results

### Variation in base composition

The base composition (frequency of the nucleotides A, C, G and T) was analysed at the three codon-positions of the *PCG-DNA* dataset (S3 Appendix). The 12 variables measured for 220 taxa were summarized by a principal component analysis (PCA) based on the first two principal components (PC), which contribute 45.80% and 24.48% of the total variance, respectively (**Figure 1**). The variables factor map shows that the variance can be explained by similar differences in base composition at the three codon-positions. Some taxa have a mtDNA genome characterized by a divergent base composition.

**Figure 1.**
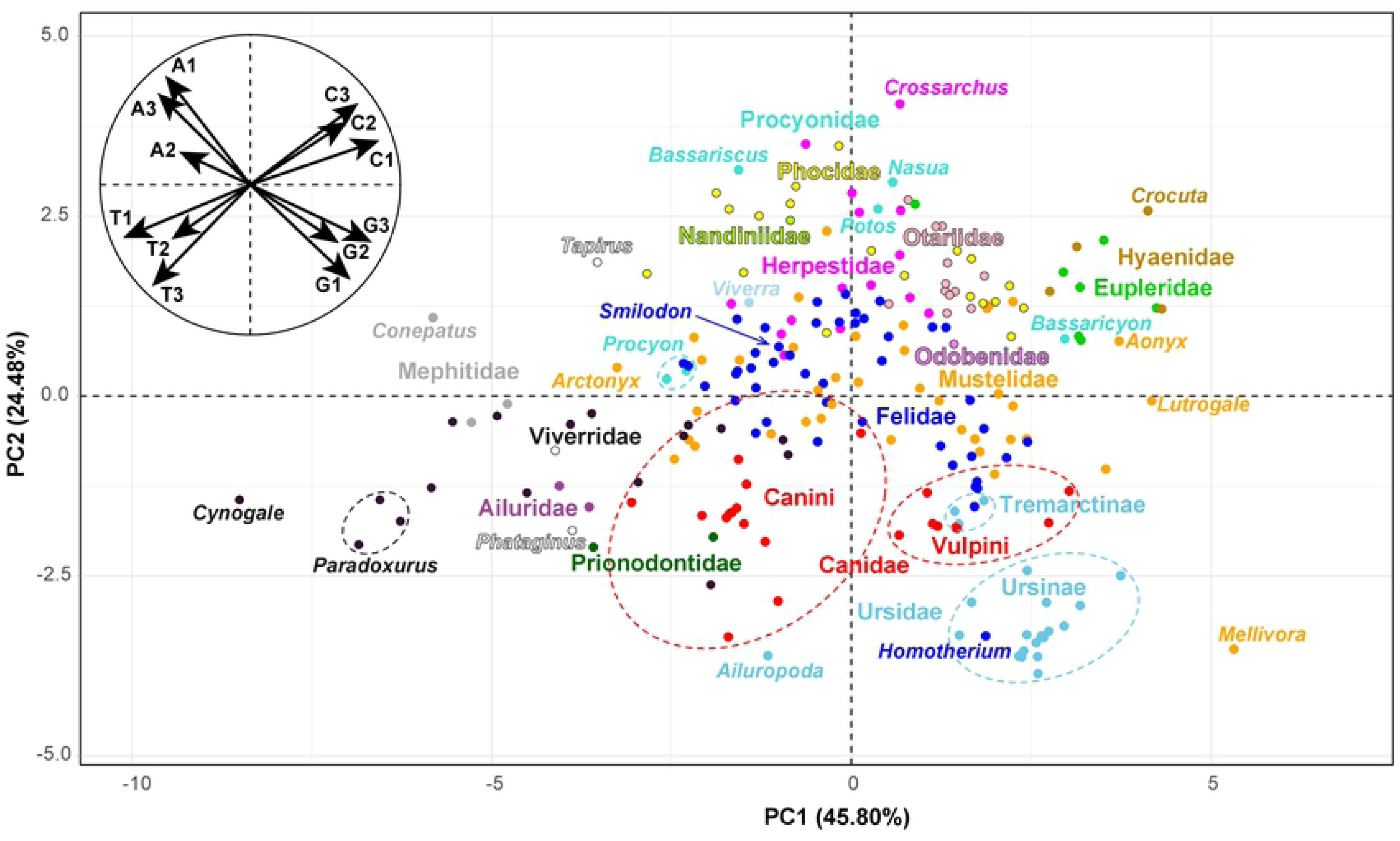
Variation in base composition of the mitogenomes of Carnivora. The *PCG-DNA* dataset was used to calculate the frequency of the four bases (A, C, G and T) at each of the three codon positions, and the 12 variables measured were then summarized by a principal component analysis (PCA). The main graph represents the individual factor map based on 220 taxa. The families of Carnivora are highlighted by different colours. The small circular graph at the top left represents the variables factor map.

Most members of the Mustelidae are found near the middle of the graph, except *Mellivora* because its mtDNA genome contains a higher percentage of guanine (15.65% *versus* “mean in other Mustelidae” [MoM] = 13.18%) and a lower percentage of adenine (28.35% *versus* MoM = 30.95%). This trend is observed at each of the three codon positions, and is more marked at third positions (S3 Appendix): G1 = 22.19% *versus* MoM = 20.87%, G2 = 12.19% *versus* MoM = 11.84%, G3 = 12.58% *versus* MoM = 6.82%; A1 = 30.55% *versus* MoM = 31.95%, A2 = 19.13% *versus* MoM = 19.41%, A3 = 35.38% *versus* MoM = 41.51%.

Within Viverridae, the mitogenome of the otter civet (*Cynogale bennettii*) is characterized by a higher percentage of thymines (32.90% *versus* “mean in other Viverridae” [MoV] = 30.73%) (**Figure 1**). This is observed at each of the three codon positions and is more marked at third positions: T1 = 25.47% *versus* MoV = 23.79%, T2 = 42.71% *versus* MoV = 42.17%, T3 = 30.52% *versus* MoV = 26.21%.

Within Ursidae, the giant panda (*Ailuropoda melanoleuca*) shows a different base composition, with more adenines (30.63% *versus* “mean in other Ursidae” [MoU] = 29.55%) and thymines (31.02% *versus* MoU = 28.98%) than other species of the family. This trend is mainly explained by differences at third codon-positions, in which the giant panda shows more adenines (40.91% *versus* MoU = 38.46%) and thymines (28.52% *versus* MoU = 23.38%).

Most members of the Felidae, including *Smilodon populator* (Machairodontinae), are found near the middle of the graph, except *Homotherium latidens* (Machairodontinae) for which the mtDNA genome shows a very atypical base composition. At third codon-positions, *Homotherium* is characterized by a lower percentage of guanine (2.80% *versus* “mean in other Felidae” [MoF] = 6.25%) and a higher percentage of thymine (25.41% *versus* MoF = 20.79%). At second codon-positions, its mitogenome contains a lower percentage of thymine (40.80% *versus* “mean in other Felidae” [MoF] = 41.89%) and a higher percentage of guanine (12.58% *versus* MoF = 11.86%). At first codon-positions, the mtDNA of *Homotherium* shows a lower percentage of adenine (30.08% *versus* “mean in other Felidae” [MoF] = 32.07%), a higher percentage of guanine (22.65% *versus* MoF = 20.92%), and a higher percentage of guanine (23.07% *versus* MoF = 22.01%).

Within Canidae, the two extant tribes can be distinguished based on their nucleotide composition, as the mitogenomes of the Vulpini (fox-like canines) show more guanines (≥ 13.89% *versus* ≤ 13.28%) and less adenines (≤ 29.53% *versus* ≥ 29.83%) than those of the Canini (dog-like canines). This trend is mainly explained by a strong bias observed at third codon-positions, in which the Vulpini taxa have more guanines (G3 ≥ 8.39% *versus* ≤ 7.30%) and less adenines (A3 ≤ 37.18% *versus* ≥ 37.80%) than the Canini.

At third codon-positions, the Eupleridae and Hyaenidae have higher percentages of cytosines than other Feliformia (mean C3: Eupleridae = 36.78%; Hyaenidae = 35.55%; other Feliformia = 30.65%).

### Pairwise mtDNA distances

Uncorrected pairwise distances were calculated on PAUP4* (Swofford, 2003) using the *mtDNA* dataset (S4 Appendix).

Intraspecific similarity distances are generally lower than 2%. However, there are several exceptions involving individuals from different geographic regions, which are or could be assigned to different subspecies or even species (see discussion): the extinct Japanese river otter (*Lutra lutra*, LC050126) *versus* extant representatives of *Lutra lutra* (2.1%); *Ursus arctos isabellinus versus* other brown bear subspecies (2.3-2.4%); *Ursus thibetanus laniger versus Ursus thibetanus mupinensis* (2.3%); the three samples of *Melogale moschata* (2.2-2.9%); the three samples of *Viverricula indica* (2.5-3.1%); *Canis lupus familiaris versus Canis lupus chanco* (2.6%); the two samples of *Paradoxurus hermaphroditus* (3.0%); the two samples of *Leopardus pardalis* (2.9%); the two mtDNA lineages identified in *Prionailurus bengalensis* (3.6%); the two samples of *Herpestes javanicus* (4.5%), although BLAST searches in NCBI suggest that the sequence NC_006835 belongs in fact to *Herpestes auropunctatus* (see discussion); the two samples of *Mungos mungo* (6.3%), but we suggest that the sample SRR7704821 may belong in fact to *Mungos gambianus* (see discussion).

Interspecific distances are generally higher than 2% of similarity. However, several species show more similar mitogenomes: *Martes martes* and *Martes zibellina* (1.9%); *Phoca largha* and *Phoca vitulina* (1.9%); *Mustela putorius versus Mustela nigripes* (1.6%) and *Mustela eversmannii* (1.2%); *Felis catus* and *Felis silvestris* (0.7%); *Zalophus californianus* and *Zalophus wollebaeki* (0.5%); *Urocyon cinereoargenteus* and *Urocyon littoralis* (0.4%). Within *Arctocephalus forsteri*, there are two mtDNA lineages: 17 mitogenomes are similar to the mitogenome of *Arctocephalus australis* (MG023139) (1.3%), whereas the 28 other mitogenomes are more divergent (2.1%). Within *Ursus arctos*, we found six mtDNA lineages showing between 1.1% and 2.4% of nucleotide divergence, but brown bears sampled on the Alaskan ABC islands (northern portion of the Alexander Archipelago) have mitogenomes which are highly similar to those of *Ursus maritimus* (0.4-0.5%).

### Phylogeny of Carnivora

The Bayesian tree of **Figure 2** was reconstructed from the *mtDNA* alignment. The results show that most nodes of the tree were highly supported (PP ≥ 0.95) and were similarly recovered as monophyletic entities using two other methods (nodes highlighted with a filled back circle in **Figure 2**), i.e. in the Bayesian tree reconstructed using the *mtDNA-Tv* alignment and JC69+I+G model (S5 Appendix) and in the Bootstrap 50% majority-rule consensus tree reconstructed from the MRP matrix of the SuperTRI analysis (S6 Appendix). Many of these nodes show maximal support in the SuperTRI analysis, as they were found monophyletic in all 10 Bayesian trees reconstructed from the 10 half-overlapping sub-datasets of the *mtDNA* alignment. For all these nodes, the phylogenetic signal is therefore robust across all parts of the *mtDNA* alignment.

**Figure 2.**
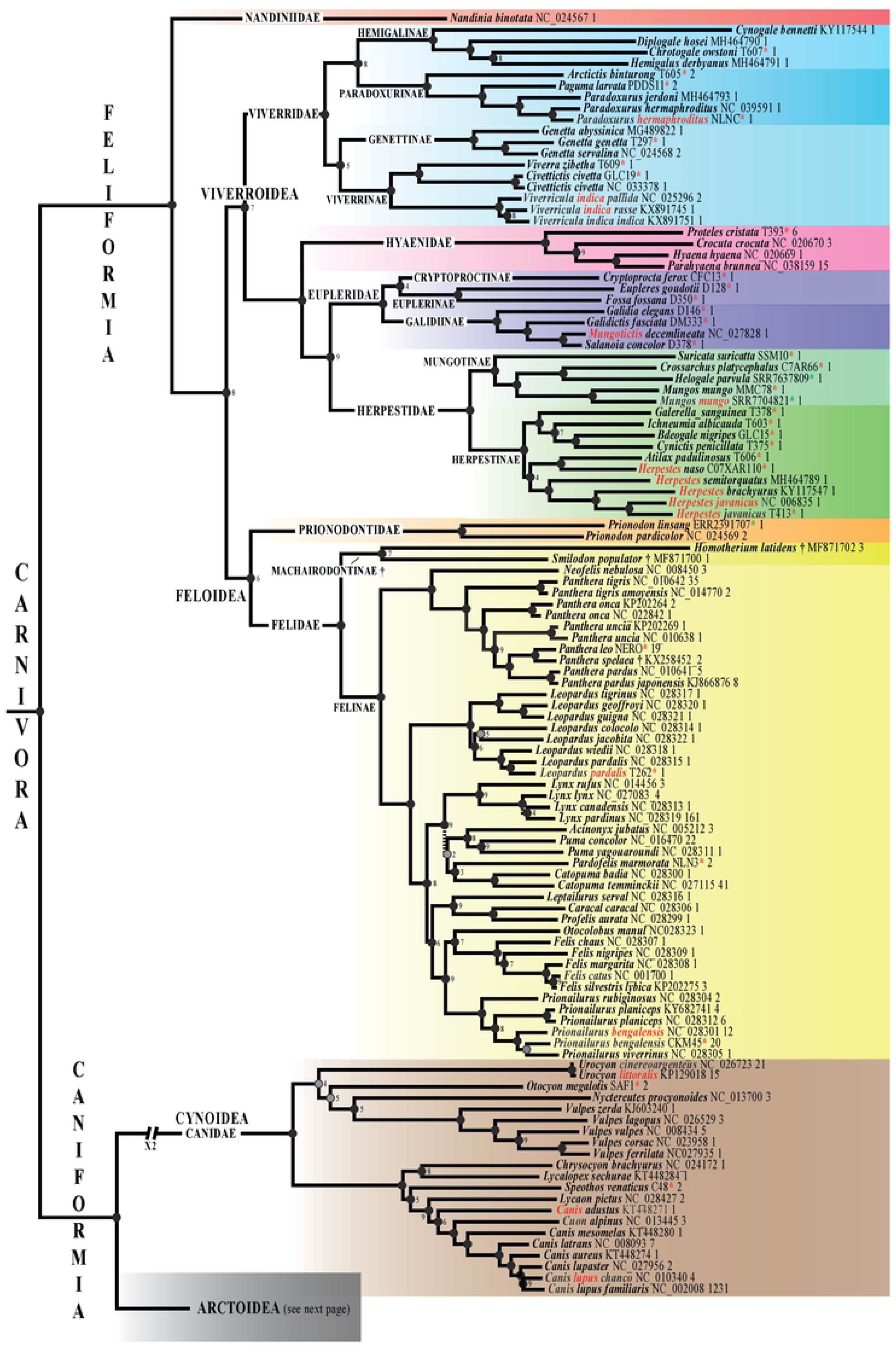
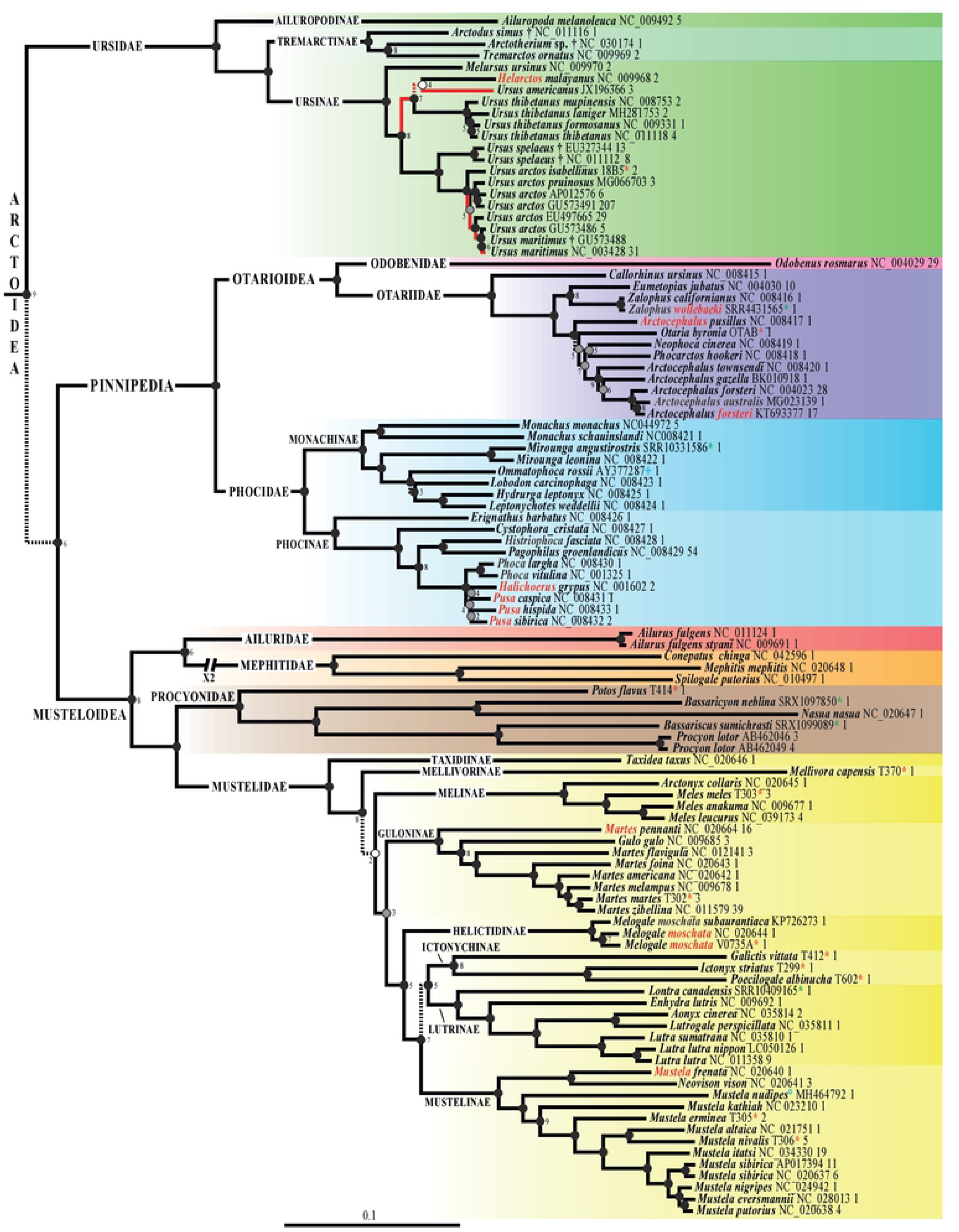
Phylogeny of Carnivora based on mitogenomes. The Bayesian tree was reconstructed using the *mtDNA* dataset (220 taxa and 14,892 bp) and GTR+I+G model. Dash branches indicate nodes supported by posterior probability (PP) < 0.95. Black circles indicate nodes that are also monophyletic in the two following trees: Bootstrap 50% majority-rule consensus tree reconstructed from the MRP matrix of the SuperTRI analysis; and Bayesian tree obtained from the analysis of the *mtDNA-Tv* dataset and JC69+I+G model. Grey circles show nodes that are not found to be monophyletic with one of the two methods detailed above. White circles indicate nodes that are not monophyletic in both *mtDNA-Tv* and SuperTRI bootstrap consensus trees. No information was provided for the nodes highly supported by the SuperTRI analysis, i.e. which were found monophyletic in all the 10 Bayesian trees reconstructed from the 10 half-overlapping sub-datasets of the *mtDNA* dataset. For nodes less supported by the SuperTRI analysis, the number of Bayesian trees (< 10) showing the nodes is indicated. Species names follow the classification of the IUCN (2020); the taxa written in red highlight the taxonomic issues discussed in the main text.

Among the nodes supported by all three methods of tree reconstruction and at least 7 of the 10 Bayesian SuperTRI analyses are the following taxa: order Carnivora; suborders Caniformia and Feliformia; infraorders Arctoidea, Cynoidea, Feloidea and Viverroidea; all the 14 families represented by at least two members in our analyses (i.e. all families except Odobenidae and Nandiniidae); the subfamilies Euplerinae, Felinae, Galidiinae, Genettinae, Guloninae, Helictidinae, Hemigalinae, Herpestinae, Ictonychinae, Lutrinae, Machairodontinae, Melinae, Monachinae, Mungotinae, Mustelinae, Paradoxurinae, Phocinae, Tremarctinae, Ursinae and Viverrinae; the genera *Catopuma*, *Felis*, *Genetta*, *Herpestes*, *Leopardus*, *Lutra*, *Lynx*, *Martes*, *Meles*, *Mirounga*, *Monachus*, *Mungos*, *Panthera*, *Paradoxurus*, *Phoca*, *Prionailurus*, *Prionodon*, *Puma*, *Urocyon*, *Vulpes* and *Zalophus*; and the species *Ailurus fulgens*, *Canis lupus*, *Civettictis civetta*, *Leopardus pardalis*, *Lutra lutra*, *Melogale moschata*, *Mustela sibirica*, *Panthera onca*, *Panthera pardus*, *Panthera tigris*, *Panthera uncia*, *Paradoxurus hermaphroditus*, *Prionailurus planiceps*, *Procyon lotor*, *Ursus maritimus, Ursus spelaeus*, *Ursus thibetanus* and *Viverricula indica.* The results support the non-monophyly of four genera: (1) *Arctocephalus*, because *Arctocephalus pusillus* is divergent from *Neophoca*, *Phocarctos* and other species of *Arctocephalus*; (2) *Canis*, because *Canis adustus* is more distantly related to other *Canis* species than to *Cuon alpinus*; (3) *Mustela*, because *Mustela frenata* is the sister-species of *Neovison vison*, whereas all other species of *Mustela* are enclosed into a robust clade; and (4) *Ursus*, because *Helarctos malayanus* is closely related to *Ursus americanus* and *Ursus thibetanus*, whereas *Ursus arctos*, *Ursus maritimus*, and *Ursus spelaeus* are grouped together. In addition, three species are not found monophyletic: (1) *Arctocephalus forsteri* is paraphyletic with respect to *Arctocephalus australis*; (2) *Prionailurus bengalensis* is paraphyletic with respect to *Prionailurus viverrinus*; and (3) *Ursus arctos* is paraphyletic with respect to *Ursus maritimus*.

### BEAST chronogram inferred from the *mtDNA* alignment

Molecular estimates of divergence times are shown in the chronogram provided in **Figure 3**. The ages were inferred with BEAST using the *mtDNA* alignment and the 22 fossil calibration points detailed in **Table 1**. Two analyses were performed using either a uniform or log-normal prior distribution on the calibrated node ages (named “U approach” and “L approach”, respectively). The two BEAST chronograms show the same tree topology (**Figure 3**), which is similar to the one reconstructed under MrBayes (**Figure 2**). As expected, the chronogram inferred with the “L approach” show more recent estimates of divergence times (values highlighted in blue in **Figure 3**) than the chronogram inferred with the “U approach”.

**Figure 3.**
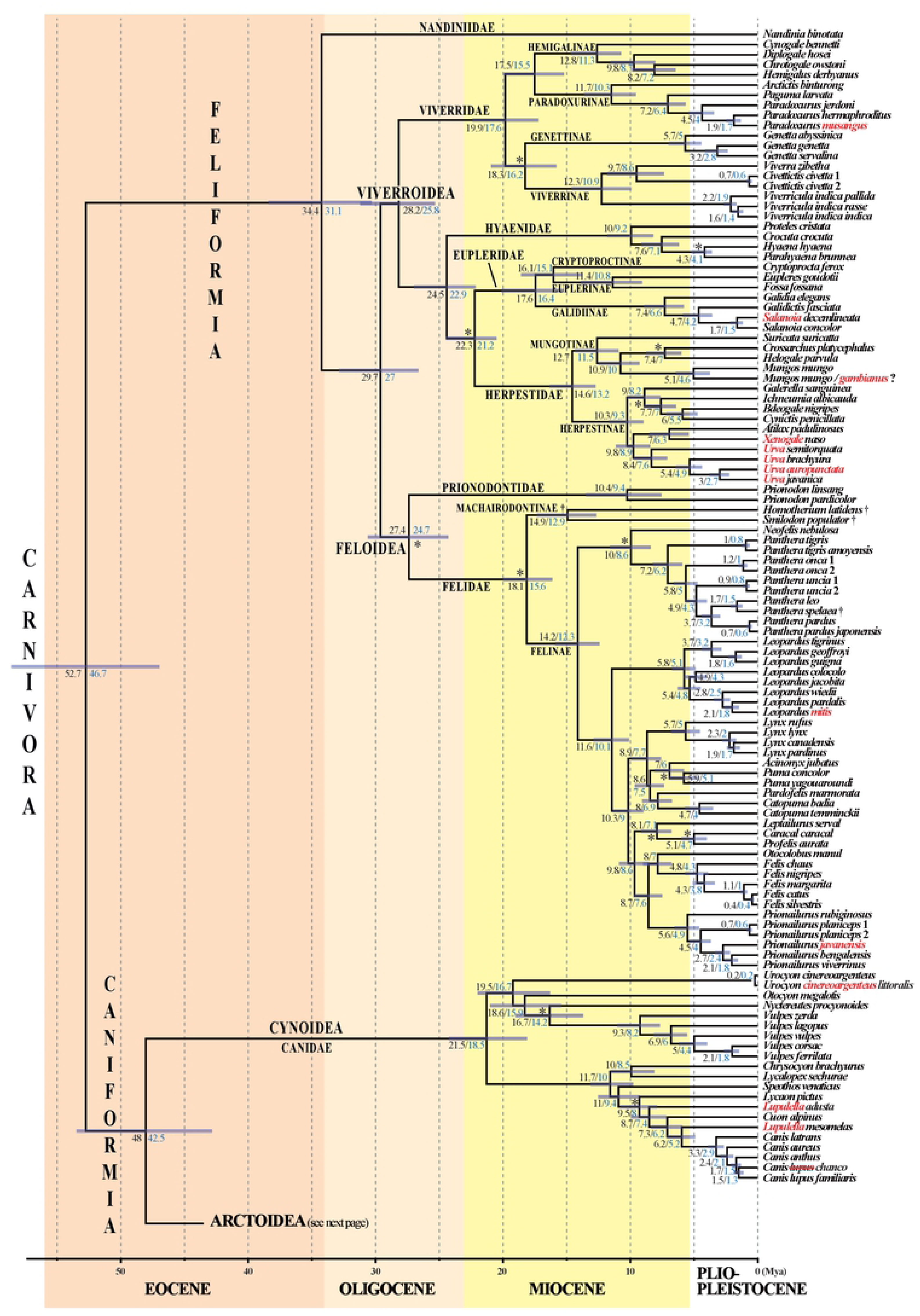
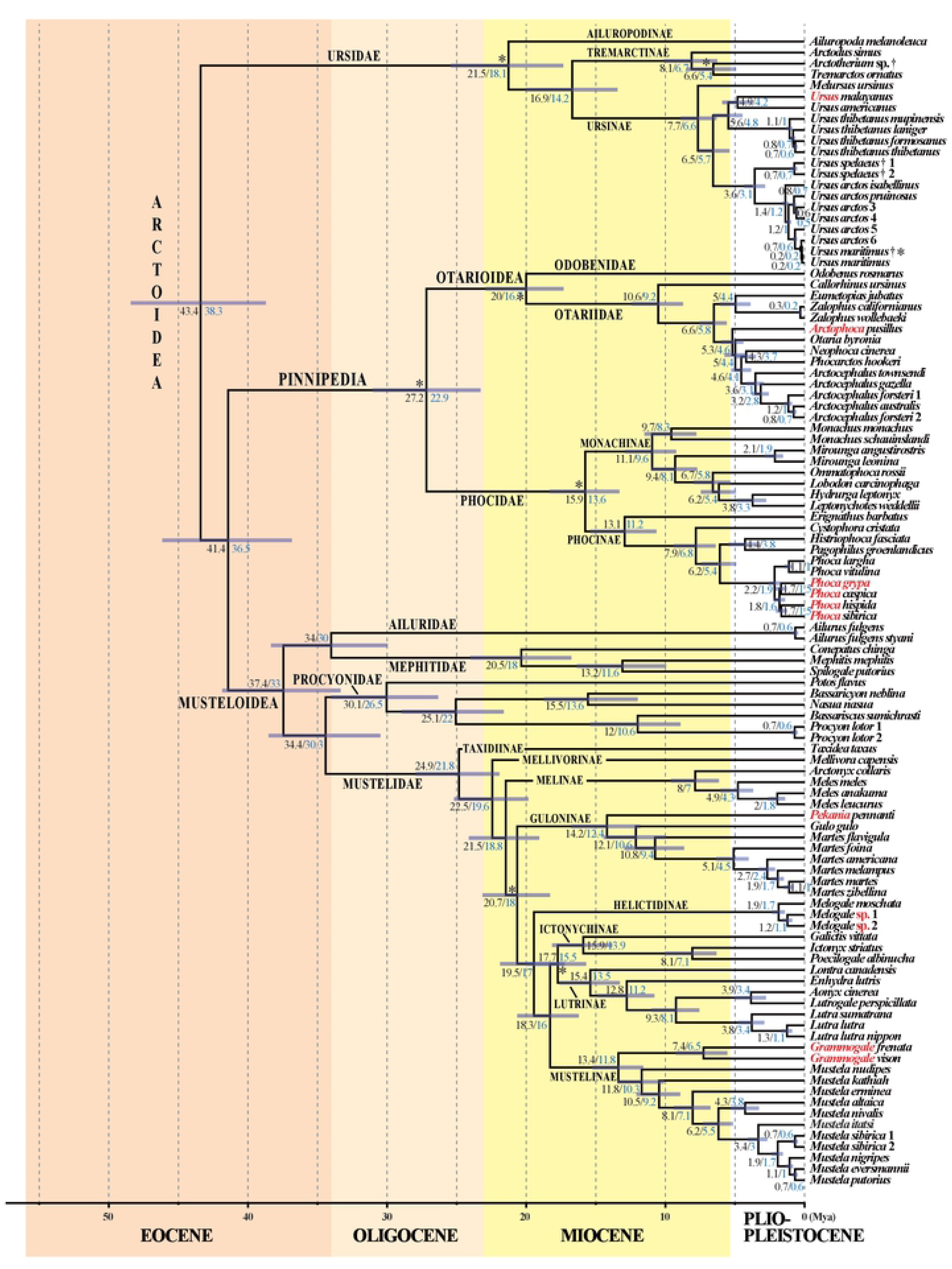
A molecular timescale for carnivoran evolution. The estimates of divergence time were calculated under BEAST v.2.4.7 using the GTR+I+G model on the *mtDNA* dataset. The asterisks show the calibration points used for molecular estimation (see Table 1 for details). The chronogram, mean ages (values in black) and associated 95% confidence intervals (grey bars) were inferred using a uniform prior distribution for fossil calibration points (“U approach”). For comparison, the values in blue are mean ages estimated using a log normal prior distribution for fossil calibration points (“L approach”; see main text for details and discussion). Species names follow the classification of the IUCN (2020); the name of the taxa written in red have been changed following our results which suggest these taxonomic changes; see the discussion for details.

Using the Geologic Time Scale v. 5.0 (Walker et al., 2018) for the correspondence between estimated divergence times and geologic epochs, the results suggest that the crown Carnivora divided into Caniformia and Feliformia during the early Eocene at around 52.7-46.7 Mya. Subsequently, the Canidae separated from other families in the early/middle Eocene at around 48.0-42.5 Mya, followed by the Ursidae and Pinnipedia in the middle/late Eocene at around 43.4-38.3 Mya and 41.4-36.5 Mya, respectively. The diversification of the Musteloidea began near the Eocene/Oligocene boundary at around 37.4-33.0 Mya, but the Mustelidae diverged from the Procyonidae in the early Oligocene at around 34.4-30 Mya. The family Phocidae separated from the Otarioidea in the Oligocene at around 27.2-22.9 Mya, whereas the separation between the Otariidae and Odobenidae occurred in the early Miocene at around 20.0-16.7 Mya. Within the Feliformia, the Nandiniidae diverged from other families at the Eocene-Oligocene transition at around 34.4-31.1 Mya. Then, the Feloidea and Viverroidea split in the Oligocene at around 29.7-27.0 Mya. The separation between the Felidae and Prionodontidae occurred in the Oligocene at around 27.4-24.7 Mya. Within the Viverroidea, the Viverridae diverged from other families in the Oligocene at around 28.2-25.8 Mya, followed by the Hyaenidae in the Late Oligocene at around 24.5-22.9 Mya, whereas the split between the Eupleridae and Herpestidae took place in the Miocene at around 22.3-21.2 Mya. Within carnivoran families, the generic diversification occurred during the Miocene (all genera of Canidae, Herpestidae, Mephitidae, Procyonidae, and Viverridae; most genera of Eupleridae, Felidae, Hyaenidae, Mustelidae, Phocidae, and Ursidae) or more rarely during the Pliocene (most genera of Otariidae, excepting *Callorhinus*; *Aonyx* / *Lutrogale*; *Caracal* / *Profelis*; *Histriophoca* / *Pagophilus*; *Hyaena* / *Parahyaena*; *Hydrurga* / *Leptonychotes*; divergence of *Galidictis*; divergence of *Helarctos*) or Pleistocene (*Halichoerus* / *Phoca* / *Pusa*; *Mungotictis* / *Salanoia*).

## Discussion

### Mitogenomic variations at the species level

In 88% of the species for which at least two mitogenomes were available (65 out of 74), intraspecific distances were found to be less than 2%. In nine species, we detected mtDNA distances greater than 2% (S4 Appendix). Such a high mitogenomic divergence suggests that the samples may belong to two or more distinct species, because of either imperfect taxonomy or species misidentification (human error). Other explanations are however possible, such as a mtDNA introgression from another species or high levels of mtDNA divergence due to strong female philopatry.

In mammals, females are generally philopatric, spending their lives close to their birthplace, whereas males typically undertake longer-distance dispersals during their lives (Greenwood, 1980; Dobson, 2013; Li & Kokko, 2019). Indeed, because they have to take care of their young, females tend to stay in areas where they can predict the risks as well as the resources. With time, this behaviour can result in a high spatial genetic structure of mtDNA variation since the mitogenome is transmitted maternally. The impact of female philopatry on the mtDNA evolution is expected to be more important if the preferred habitat-type is delimited by strong geographic barriers representing a high risk for the dispersal of females (and their young), for example, open areas for small forest fruit bats (Hassanin et al., 2015) or large rivers and mountains for giraffes (Petzold & Hassanin, 2020). Female philopatry may therefore result in the selection of highly divergent mtDNA haplotypes in geographically isolated maternal lineages, whereas gene flow can be still maintained through long-distance male dispersals. Female philopatry can be advanced to explain the high intraspecific mtDNA distances found between populations of bears of the species *Ursus arctos* (6 lineages, 1.1-2.4%) and *Ursus thibetanus* (4 linages, 1.0-2.3%). This hypothesis is supported by previous phylogeographic studies showing strong geographic structure of mtDNA variation (Hirata et al., 2013; Wu et al., 2015) and by field data on brown bears indicating that natal dispersal distances are five times smaller for females (Steyaert et al., 2012).

In several other species, we found two or three mitogenomic haplotypes or haplogroups which are separated by more than 2%. In the Eurasian otter (*Lutra lutra*), Waku et al. (2016) sequenced the mtDNA genome of three extinct river otters of Japan: the two mtDNA genomes of otters from Honshu are similar to those found in extant Eurasian otters (< 1%), whereas the genome of the otter from Shikoku Island differs by 2.1%, suggesting that it belongs to a distinct species (*Lutra nippon*) or subspecies (*Lutra lutra nippon*). Since female philopatry *versus* male biased dispersal was well attested by both radiotracking and genetic data in European otters (Quaglietta et al., 2013), we consider that nuclear data are needed to solve this taxonomic issue.

The three mtDNA haplotypes available for the small-toothed ferret badger (*Melogale moschata*) show comparatively high nucleotide distances, i.e., between 2.2 % and 2.9 %. The genus *Melogale* includes five species (IUCN, 2020): *Melogale cucphuongensis* known from a few animals collected in northern Vietnam (Nadler et al., 2011; Li et al., 2018), *Melogale everetti* on Borneo, *Melogale moschata* which is widely distributed from North-east India to Taiwan through China and Indochina, *Melogale orientalis* on Java, and *Melogale personata*, which is found in mainland Southeast Asia. In agreement with the recent study of Rozhnov et al. (2019), our phylogenetic analysis based on cytochrome b (*CYB*) sequences (S7 Appendix) shows that *Melogale cucphuongensis* is the sister-group of a clade composed of three divergent mtDNA lineages: (i) *Melogale personata*, as represented by 16 samples from Vietnam; (ii) a first mtDNA haplogroup of *Melogale moschata*, which includes five samples from Vietnam (including our V0735A sample) and the reference mitogenome (NC_020644, unknown origin); and (iii) a second mtDNA haplogroup of *Melogale moschata*, which contains a sample from Taiwan (subspecies *subaurantiaca*) and a sample from China (MN400429, from Fei Zhou). Since the holotype of *Melogale moschata* was collected in the Guangdong Province of South China (Storz & Wozencraft, 1999), our *CYB* tree suggests that the two samples from China and Taiwan represent the species *Melogale moschata*, whereas the samples from Vietnam currently assigned to *Melogale moschata* may belong to a new species, confirming that Vietnam is a key hot spot for the genus *Melogale*.

Three mitogenomic haplotypes were available for the small Indian civet (*Viverricula indica*), representing the three main mtDNA lineages previously identified in Gaubert et al. (2017), i.e. the subspecies *V. i. indica* from India, Madagascar (where it was introduced), Sri Lanka, West China and northern Indochina; *V. i. rasse* from southern Indochina to Java; and *V. i. pallida*, from East China to Taiwan. The mitogenomic divergence are between 2.5% and 3.1%, suggesting that they may be treated as different species, but nuclear markers are needed to further investigate this taxonomic issue. Gaubert et al. (2017) concluded that these three mitogenomic lineages diverged from each other in the Pliocene, between 3.2 Mya and 2.6 Mya, but our dating estimates rather support that the divergences occurred in the Pleistocene, between 2.2-1.9 Mya and 1.6-1.4 Mya.

In the common palm civet (*Paradoxurus hermaphroditus*), the two mtDNA haplotypes differ by 2.9%, suggesting that they may come from two separate species. The phylogeographic studies of Patou et al. (2010) and Veron et al. (2015) based on both mitochondrial and nuclear sequences (*CYB*, control region, intron 7 of *FGB*) supported the existence of three distinct species of *Paradoxurus*: *P. hermaphroditus* (Indian and Indochinese regions), *P. musangus* (mainland Southeast Asia, Sumatra, Java and other small Indonesian islands) and *P. philippinensis* (Mentawai Islands, Borneo and the Philippines). Only the first two of them were sampled in our mitogenomic study. Since they have overlapping range in mainland Southeast Asia, where it can be assumed that *P. hermaphroditus* dispersed from the north of India while *P. musangus* arrived from the south (see Figure 2 in Patou et al., 2010), the possibility of interspecific hybridization should be investigated in this region.

The two mitogenomes sequenced for the ocelot (*Leopardus pardalis*) differ by 3.0%, suggesting that this taxon may be split into two separate species, as proposed by Nascimento (2010) based on a morphological analysis of 591 specimens of *Leopardus* spp. Using an alignment of the 5’-part of the mitochondrial control region, Eizirik et al. (1998) have suggested a further division into four geographic groups. However, there is a 100-bp stretch of missing data in all their sequences, which is located in a region where we detected three motifs of 80 bp repeated in tandem in the mitogenome of *Leopardus pardalis mitis* and only two motifs in that of *Leopardus pardalis pardalis*. These repetitive sequences, named RS2 in Hoelzel et al. (1994), were found in variable number in all families of Feliformia, from two in *Prionodon pardicolor* (Hassanin, 2016) to five in *Nandinia binotata* (Hassanin & Veron, 2016). These variations in RS2 tandem repeats may pose serious problems of homology for DNA alignment. If RS2 repeats are removed from the alignment of Eizirik et al. (1998), there is no robust signal for phylogenetic relationships within *Leopardus pardalis* (data not shown). We recommend therefore to study the phylogeography of ocelot lineages with mitochondrial protein-coding genes, such as *CYB* and *CO1* genes, or full mitogenomes. For taxonomy purpose, the analyses should be completed with nuclear markers, as Wultsch et al. (2016) have shown evidence for female philopatry *versus* male-biased dispersal in ocelots sampled in Belize (Central America).

Two divergent mtDNA haplogroups were found for the leopard cat *Prionailurus bengalensis* (3.6%), corresponding to the Asian mainland leopard cat and Sunda leopard cat (Luo et al., 2014; Patel et al., 2017). A species-level separation was confirmed by Y chromosome and whole-genome SNP studies (Luo et al., 2014; Li et al., 2016). Therefore, we follow Kitchener et al. (2017) in recognizing two species of leopard cats: the mainland leopard cat *Prionailurus bengalensis* for Asian mainland leopard cats, and the Sunda leopard cat *Prionailurus javanensis* for leopard cats from Java, Sumatra and Borneo. Previous molecular estimates have provided different ages for their divergence, between 2.67 Mya and 0.92 Mya (Luo et al., 2014; Li et al., 2016; Patel et al., 2017). Although based on different calibration points, our estimate of 2.7-2.4 Mya corroborates that of Luo et al. (2014). This suggests that maternal linages were first isolated at the Pliocene/Pleistocene boundary, when the glacial/interglacial oscillations started. However, the mitogenomic tree does not support a sister-group relationship between the two species of leopard cats, because *Prionailurus bengalensis* is closer to *Prionailurus viverrinus*, the fishing cat, than to *Prionailurus javanensis*. This result contrasts with the nuDNA trees of Luo et al. (2014) and Li et al. (2016) showing a sister relationship of *Prionailurus bengalensis* and *Prionailurus javanensis*. In agreement with the scenario proposed by Li et al. (2016), such a discordance suggests that a mtDNA introgression occurred from *Prionailurus bengalensis* to *Prionailurus viverrinus* in the Pleistocene epoch, here dated at 2.1-1.8 Mya.

The two mitogenomes of the Javan mongoose (*Urva javanica*, previously named *Herpestes javanicus*, see Patou et al., 2009 for details) differ by 4.5% in our alignment. BLAST searches in NCBI show that the reference genome sequenced for *Urva javanica* (NC_006835) belongs in fact to its sister-species, *Urva auropunctata* (Small Indian mongoose, previously named *Herpestes javanicus* or *Herpestes auropunctatus* (Patou et al., 2009). In agreement with this, the sample was collected in Fiji where the Small Indian mongoose is known to have been introduced (Veron & Jennings, 2017).

Two highly divergent mtDNA haplotypes (6.3%) were detected in banded mongooses *Mungos mungo* from Halle Zoological Garden (Germany) and from the San Diego Zoo Institute for Conservation Research (USA). The geographic origins of these animals are unknown, but such a high pairwise distance suggests that the two mitogenomes may represent the two species currently described in the genus *Mungos*: the banded mongoose (*Mungos mungo*), which is widely distributed in sub-Saharan Africa, and the Gambian mongoose (*Mungos gambianus*), which is endemic to West Africa. BLAST searches in NCBI show that our MMC78 sample share 99.9% of *CYB* identity with a mongoose from Kenya (AY928674; Koepfli et al., 2006), confirming that it belongs to the species *Mungos mungo*. By contrast, the sample SRR7704821 shows only 93% of identity with *Mungos mungo* AY928674, suggesting that alternatively it may belong to *Mungos gambianus*. Morphologically, the two species are similar but *Mungos mungo* is characterized by 10 to 15 dark dorsal stripes that are absent in *Mungos gambianus*. These marked differences render the hypothesis of misidentification unlikely. As an alternative, a mitochondrial introgression from *Mungos gambianus* to *Mungos mungo* may have occurred either in captivity or in the wild, as the two species can be found in sympatry in West Africa (IUCN, 2020).

### Shallow mitochondrial phylogeny

The mtDNA distances calculated between closely-related species are generally higher than 2% (73% of the comparisons detailed in S4 Appendix). However, there are several exceptions in the genera *Arctocephalus*, *Felis*, *Martes*, *Mustela*, *Phoca*, *Urocyon*, *Ursus* and *Zalophus*. Four main hypotheses can be advanced to explain the existence of similar mtDNA genomes in two putative species: species misidentification (human error), imperfect taxonomy (species synonymy), mtDNA introgression, or recent speciation event during the Pleistocene (Hassanin et al., 2012).

Within *Arctocephalus*, two divergent mtDNA lineages (2.1%) were sequenced for the New Zealand fur seal (*Arctocephalus forsteri*) (Emami-Khoyi et al., 2017). One of them was found to be more similar to the single mitogenome available for the South American fur seal, *Arctocephalus australis* (1.3%). The mitochondrial paraphyly of *Arctocephalus forsteri* suggests a mtDNA introgression from *Arctocephalus australis* at 0.8-0.7 Mya. Nuclear data should be sequenced to confirm this hypothesis and to further explore a possible taxonomic issue between South American and New Zealand fur seals.

Within *Felis*, the mitogenome of the domestic cat (*Felis catus*) is very similar to those of the wild cat, *Felis silvestris* (0.7%), including the African wild cat (*Felis silvestris lybica*), European wild cat (*Felis silvestris silvestris*) and Chinese desert cat (*Felis silvestris bieti*). These three subspecies and the domestic cat are either considered separate species (Kitchener et al., 2017) or subspecies of *Felis silvestris* (e.g. Driscoll et al., 2007; Mattucci et al., 2015). Such a genome similarity between the domestic cat and wild cat was expected because the cat was domesticated from the near Eastern wild cat *Felis silvestris lybica* (Driscoll et al., 2007) and can hybridize with wild forms (Randi, 2008). Our results confirm that these taxa should be considered subspecies rather than species.

Within *Martes*, the Pine marten (*Martes martes*) from western Europe and the Sable (*Martes zibellina*) from Siberia and adjacent areas have quite similar mitogenomes (1.9% divergence), and our estimation of divergence time is 1.1-1.0 Mya, which is in perfect agreement with Law et al. (2018). However, Rozhnov et al. (2010) have detected high levels of reciprocal mtDNA introgression in northern Urals, where populations of the two species are found in sympatry. In this hybrid zone, levels of gene flow should be further studied using nuclear markers to determine if introgression between the two taxa is asymmetric or not.

Within *Mustela*, several species show low levels of mitogenomic divergence (1.2-1-6%), including the western polecat (*M. putorius*) distributed in western Europe, the steppe polecat (*M. eversmanii*) found in eastern Europe and Asia, and the black-footed ferret (*M. nigripes*) endemic to North America. In addition, the mitogenome of the European mink, *M. lutreola* (MT304869, not included in our phylogenetic analyses) shares between 99.1% and 99.2 % of identity with the four mitogenomes available for *M. putorius*. Our estimations indicate that these four closely related species of *Mustela* have diverged from each other after 1.1-1.0 Mya. However, *M. putorius*, *M. eversmanii*, and *M. lutreola* can be found in sympatry in Europe (IUCN, 2020), suggesting possible mtDNA introgression during the Pleistocene epoch. Cabria et al. (2011) have shown evidence for mtDNA introgression from *M. lutreola* to *M. putorius*, resulting apparently from asymmetric interspecific hybridization, i.e. involving only females of European mink and males of polecat. Since the western and steppe polecats are known to occasionally hybridize in Eastern Europe (Abramov et al., 2016), gene flow should be also studied between the two taxa.

Within *Phoca*, harbor seals (*P. vitulina*) and spotted seals (*P. largha*) are parapatric sibling species, showing 1.9% of differences in our mitogenomic alignment. Although the two species are known to hybridize in captivity (IUCN, 2020), they were found reciprocally monophyletic in previous mtDNA studies (Nakagawa et al., 2010; Cordes et al., 2017) and there is no evidence of mixed ancestry in wild populations found in sympatry (e.g. in Alaska; Cordes et al., 2017). The hypothesis of a recent Pleistocene speciation, at 1.1-1.0 Mya according to our estimation, is therefore the most likely.

Within *Urocyon*, the island fox (*U. littoralis*), restricted to six California Channel Islands located off the coast of southern California, USA, and the mainland grey fox (*U. cinereoargenteus*) share very similar mtDNA genomes (0.4%). In addition, the species *U. cinereoargenteus* was found to be paraphyletic in the mitochondrial tree of Hofman et al. (2015) due to the inclusive placement of *U. littoralis*. Even if the island fox is approximately 25% smaller than the mainland grey fox (Schutz et al., 2009), we suggest that the island fox should be rather treated as a subspecies of *U. cinereoargenteus*. Nuclear genome comparisons are however needed to definitively clarify this taxonomic issue.

Within *Ursus*, bears living on the Alaskan ABC islands have mitogenomes which are very similar (0.4-0.5%) to those sequenced for extant polar bears and the fossil of Svalbard dated between 130 ka and 110 kya (Lindqvist et al., 2010). Two possible scenarios of mitochondrial introgression have been previously proposed: Hailer et al. (2012) suggested that the ancestor of polar bears was introgressed by brown bears between 166 and 111 kya; whereas Hassanin (2015) suggested that different populations of brown bears were introgressed by polar bears at two glacial periods of the Pleistocene, at 340 ± 10 ka in western Europe, and at 155 ± 5 ka on the ABC islands, and probably also in Beringia and Ireland based on ancient DNA sequences.

Within *Zalophus*, two species are currently recognized: the Californian Sea Lion (*Z. californianus*), which is found on the Pacific coasts of North America; and the Galápagos sea lion (*Z. wollebaeki*), which has been considered as a subspecies of *Z. californianus* by some authors (Aurioles-Gamboa & Hernández-Camacho, 2015). Despite the low mitogenomic distance measured here between the two species (0.5%), Wolf et al. (2007) have previously concluded that they are separate species because they were found reciprocally monophyletic with D-loop and cytochrome b sequences and numerous private alleles were detected for both taxa at most of the 25 investigated microsatellite loci. Our molecular estimate of divergence time between *Z. californianus* and *Z. wollebaeki* is 0.3-0.2 Mya, which is much more recent than previous estimations based on D-loop sequences, i.e. 0.8 Mya (Schramm et al., 2009) and 2.3 ± 0.5 Ma (Wolf et al., 2007).

### Mitogenomic variations at the genus level

In Mammals, most events of interspecific diversification at the genus level are more recent than 10 Mya and the great majority of them occurred during the Pliocene and Pleistocene epochs. This trend, previously reported in various genera of Cetartiodactyla (Hassanin et al., 2012) and Primates (Pozzi et al., 2014), is here confirmed in the following 25 carnivoran genera: *Arctocephalus*, *Catopuma*, *Felis*, *Genetta*, *Leopardus*, *Lutra*, *Lynx*, *Martes*, *Meles*, *Melogale*, *Monachus*, *Mirounga*, *Mungos*, *Mustela*, *Panthera*, *Paradoxurus*, *Phoca*, *Prionailurus*, *Prionodon*, *Puma*, *Pusa*, *Ursus*, *Urva* and *Vulpes*. Our analyses suggest however that two mustelid species separated from their congeneric representatives in the Middle Miocene: *Martes pennanti* at 14.2-12.4 Mya; and *Mustela frenata* at 13.4-11.8 Mya. In addition, the phylogenetic positions of these two species result in the polyphyly of the two genera *Martes* and *Mustela*, as previously found using both mtDNA and nuDNA markers (Law et al., 2018). Taken together, these results clearly indicate that *Martes pennanti* should be placed in its own genus *Pekania*, whereas *Mustela frenata* should be placed in the genus *Grammogale*. Indeed, the genus *Grammogale* Cabrera, 1940 was used by Cabrera & Yepes (1960) to unite *Mustela africana* and *Mustela felipei* into a separate genus. In addition, several molecular studies have shown that these two species fall into a robust clade with *Mustela frenata* and that they are the sister-group of *Neovison vison* (Harding & Smith, 2009; Law et al., 2018). We recommend therefore to include the four species *M. africana*, *M. felipei*, *M. frenata*, and *N. vison* into the genus *Grammogale*.

Within the tribe Canini, our mitogenomic tree shows that *Cuon alpinus* (dhole) is the sister-group of all *Canis* species, excepting *Canis adustus* (side-striped jackal) which diverged from them at 8.7-7.4 Mya. The African wild dog (*Lycaon pictus*) occupies a more basal position, but this placement is only supported by 6/10 overlapping datasets in our SuperTRI analyses, indicating that an alternative position cannot be excluded. The mitochondrial phylogeny differs from the nuclear phylogeny of Lindblad-Toh et al. (2005) and Koepfli et al. (2015) in which *C. adustus* was found to be related to *Canis mesomelas* (black-backed jackal), but more divergent from the clade composed of other *Canis* species, *Cuon* and *Lycaon*. Following these results, in order to keep the genus *Canis* monophyletic, the species *C. adustus* and *C. mesomelas* should be placed in a different genus, which is *Lupulella* according to Viranta et al. (2017). The mito-nuclear discordance for the monophyly of the genus *Lupulella* was not discussed in previous studies. Our interpretation is that a mitochondrial introgression occurred at 6.2-5.2 Mya from an ancestor of *Canis* species to the lineage leading to *Lupulella mesomelas*. In agreement with the genomic study of Gopalakrishnan et al. (2019), which concluded to pervasive gene flow among *Canis* species, our results suggest that interspecies hybridization has been also frequent in the early evolutionary history of canid genera in Africa.

Our chronogram in **Figure 3** shows that no intergeneric divergence occurred during the Pleistocene epoch, except the split between the Malagasy genera *Mungotictis* and *Salanoia*, and the separation between the seal genera *Halichoerus*, *Phoca* and *Pusa*. A Pleistocene diversification was also found in previous molecular dating analyses on the Eupleridae and Phocidae (Higdon et al., 2007; Nyakatura & Bininda-Emonds, 2012). In the absence of striking morphological feature for diagnosing these genera, we recommend synonymizing *Halichoerus* and *Pusa* with *Phoca*, as proposed in previous studies (e.g. Arnason et al., 2006), and *Mungotictis* with *Salanoia*.

### Changes in base composition and their impact on evolutionary rates

Mitogenomic rearrangements did not occur during the evolutionary history of the Carnivora. This deeply contrasts with other animal groups for which inversions of mitochondrial protein-coding genes and control region have resulted in many local or full reversals of base composition (see Hassanin et al., 2005; Arabi et al., 2010 for more details). Despite this, five carnivoran species in our Bayesian tree (**Figure 2**) have a longer terminal branch than other representatives of the same family or superfamily: *Cynogale bennetti* within the Viverridae, *Homotherium latidens* within the Felidae, *Mellivora capensis* within the Mustelidae, *Nasua nasua* within the Procyonidae, and *Odobenus rosmarus* within the Otarioidea. In addition, two higher taxa of Caniformia show longer branches: Otarioidea and Musteloidea. Since branch length is proportional to the number of substitutions, these long branches indicate either an acceleration in the rates of substitution or alternatively an important change in the pattern of substitution (Tamura & Kumar, 2002; Hassanin, 2006). As shown in **Figure 1**, *Cynogale*, *Homotherium*, and *Mellivora* show a divergent base composition. For *Cynogale* and *Mellivora*, we can therefore assume that a change in the pattern of substitution has taken place during their recent evolution. The case of *Homotherium* is more problematic. The mitogenome of this extinct genus was sequenced from a bone (YG 439.38) dated at >47,500 years (Paijmans et al., 2017). Since ancient DNA molecules are known to exhibit a high rate of cytosine deamination, the sequences can contain artefactual C=>T and G=>A substitutions (Briggs et al., 2010). By comparing the mitogenomes of all felid species, this is exactly the pattern observed for third-codon positions, as *Homotherium* is characterized by less cytosines (28.94% *versus* 28.96-35.21% in other felid species), more thymines (25.41% *versus* 17.44-23.51%), less guanines (2.80% *versus* 4.69-8.41%) and more adenines (42.85% *versus* 38.41-42.79%). As a consequence, we conclude that sequencing errors introduced by DNA damage are the cause of the long branch of *Homotherium* in the tree of **Figure 2**.

The mitogenomes of *Nasua* and *Odobenus* have a base composition that is not so divergent from their closest relatives, suggesting that an acceleration of rates of substitution is the cause of their long branch in the tree of **Figure 2**. As pointed out by Bromham (2011), mammals evolving with faster evolutionary rates have generally a small body-size, but the cause of this body-size effect continues to be debated, because there are many possible mechanisms including shorter generation times, shorter lifespan, higher fecundity, larger litter size, and higher metabolic rates. From this point, the walrus (*O. rosmarus*) is intriguing because it is one of largest pinnipeds. Its field metabolic rate is 381.2 MJ/day, which is 6 to 32 times more than in other species of Pinnipedia (Acquarone et al., 2006). Whereas most other pinnipeds hunt pelagic organisms, such as fish and cephalopod species, in the water column, the walrus feeds on benthic species and prefers molluscs, especially clams (Lowry, 2016). This special diet and associated behaviour, with long exposure in the cold coastal waters of the Arctic Ocean, may therefore explain the increased rate of mtDNA evolution detected in the walrus. Repetitive bottlenecks generated by climatic oscillations during the Pleistocene may be proposed as an alternative hypothesis. Nuclear genome comparisons between pinniped species will help to decipher between these two main hypotheses, as we can expect accelerated rates of substitution in both mtDNA and nuDNA genomes in case of repetitive bottlenecks.

### Robust and reliable mitogenomic phylogeny for deep relationships within Carnivora

Previous molecular studies have shown that inferring deep phylogenetic relationships using mitogenomes can be problematic. An important issue concerns the reversals of strand mutational bias: detected in many animal phyla, such as Arthropoda, Chordata, Mollusca, etc, it may be particularly misleading for phylogenetic reconstruction (Hassanin et al., 2005; Arabi et al., 2010). The mitogenome of mammals is not affected by this kind of bias, as its structure is highly conserved among the 27 mammalian orders. However, mutational saturation is a different issue: since the mtDNA evolves with higher mutation rates than the nuclear genome (Brown et al., 1979), it is more prone to multiple substitutions at the same site which leads, with time, to the disappearance of the phylogenetic signal (Philippe & Forterre, 1999; Xia et al., 2003). Although mutational saturation may be particularly problematic for reconstructing Precambrian, Paleozoic, and Mesozoic divergences, it is expected to have less impact for inferring more recent divergences, such as those during the Cenozoic. This explains why mitogenomic sequences have been largely used for inferring interfamilial and intrafamilial relationships in most mammalian groups, including Cetartiodactyla (Hassanin et al., 2012), Feliformia (Zhou et al., 2017), Primates (Finstermeier et al., 2013; Pozzi et al., 2014), Pholidota (Hassanin et al., 2015), Xenarthra (Gibb et al., 2016), etc. In agreement with this view, all families and inter-familial levels shown in our mtDNA tree of **Figure 2** received high support with all the three phylogenetic approaches (Bayesian trees of *mtDNA* and *mtDNA-Tv* data sets, and SuperTRI analysis), indicating considerable stability of the topology. Interfamilial relationships are conformed to the nuclear tree published by Eizirik et al. (2010), except the position of the Mephitidae as the sister-group of the Ailuridae. However, this relationship was also found by Law et al. (2018) based on 46 genes (4 mitochondrial and 42 nuclear genes) and 75 species of Musteloidea.

### Diversification of the Carnivora during the Cenozoic

The mtDNA phylogeny reconstructed here presents a good opportunity to study the evolution of the order Carnivora. Indeed, our dense taxonomic sampling allow us to include many fossils as calibration points for estimating divergence times. This point is particularly relevant as the fossil record of Carnivora has been significantly improved over the last ten years, with the discovery of several key fossils (e.g., Abella et al., 2012; Grohé et al., 2013; Tseng et al., 2014; Berta et al., 2018; Valenciano et al., 2020). As pointed out in Warnock et al. (2015), the most effective means of establishing the quality of fossil-based calibrations is through a priori evaluation of the intrinsic palaeontological, stratigraphic, geochronological and phylogenetic data. We identified therefore 22 calibration points (**Table 1**), including one tip calibration (the mitogenome of a late Pleistocene fossil of *Ursus maritimus* published in Lindqvist et al., 2010) and 20 well-constrained calibration points having both minimum and maximum age boundaries interpreted from fossils with known position with respect to extant taxa included in our study. Molecular dating analyses were performed using either a uniform (U) or log-normal (L) prior distribution on the calibrated node ages. The “L approach” used here considers that minimum ages are generally more accurate and reliable than maximum ages because younger fossils are generally more abundant and precisely dated than older fossils as a consequence of taphonomy and dating methods (Crees et al., 2019). As expected, the divergence times estimated with the “L approach” are more recent than those estimated with the “U approach” (**Figures 3 and 4**). The values calculated with the two approaches are given for all the nodes described below.

**Figure 4.**
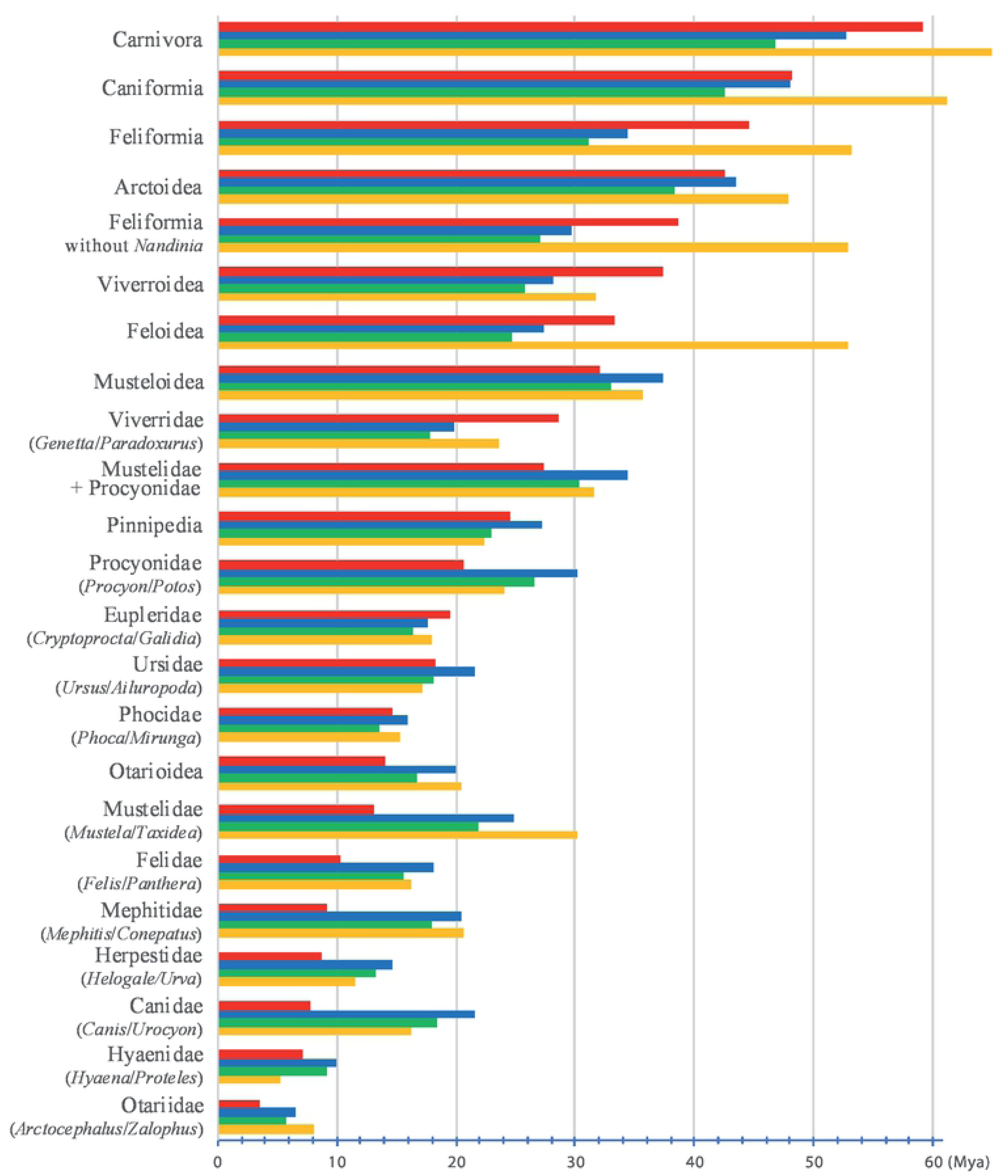
Comparison with published chronograms on Carnivora. The mean divergence times were here estimated with two approaches for the prior distribution on the calibrated node ages: (1) a uniform distribution between maximum and minimum boundaries (“U approach”, blue histograms); and (2) a log-normal distribution (“L approach”, green histograms; see main text for details). The results were compared with mean ages inferred in Eizirik et al. (2010; red histograms) and Nyakatura & Bininda-Emonds (2012; orange histograms).

The age estimated for the MRCA of Carnivora was 52.7 Mya with the “U approach” and 46.7 Mya with the “L approach”, and both are younger than previous estimates published in Eizirik et al. (2010; 59.2 Mya) and Nyakatura & Bininda-Emonds (2012; 64.9 Mya). However, our dating estimates fit well with the end of the warmest period of the Cenozoic era, from the Palaeocene–Eocene Thermal Maximum (PETM) at 56 Mya to the Early Eocene Climatic Optimum (EECO) at around 53-50 Mya (Zachos et al., 2001; Zachos et al., 2008; Kirtland Turner et al., 2014). In the fossil record, this period was marked by the first appearances of Primates, Perissodactyla and Cetartiodactyla in North America and Europe, as well as Carnivoraforms, a group formed by the crown group Carnivora plus the stem family ‘Miacidae’ (Spaulding and Flynn, 2012; Hooker 2015; Solé et al., 2016). During the early Eocene greenhouse world, rainforests spread on Earth from pole to pole (Sluijs et al., 2006; Pross et al., 2012), suggesting that early carnivorans were small arboreal species. Biogeographically, Carnivoraforms diversified in the three Laurasian continents during the early Eocene, with a possible origin in the late Paleocene of Asia (Solé et al., 2016). Crown carnivorans originated in one of the three continents of the Northern Hemisphere. To know which one, new fossils need to be discovered in the middle Eocene to fill the gap between the Carnivoraforms of the early Eocene and the oldest carnivorans of the late Eocene (Solé et al., 2016).

The divergence times estimated for basal relationships in the Caniformia are similar to those published in Eizirik et al. (2010) but much younger than those published in Nyakatura & Bininda-Emonds (2012): Caniformia = 48.0-42.5 Mya *versus* 48.2 Mya and 61.2 Mya respectively; Arctoidea = 43.4-38.3 Mya *versus* 42.6 Mya and 47.8 Mya. For more recent nodes, our estimates are similar to the two previous studies: Musteloidea = 37.4-33.0 Mya *versus* 32.0 Mya and 35.7 Mya; Mustelidae+Procyonidae = 34.4-30.3 Mya *versus* 27.4 Mya and 31.6 Mya; and Pinnipedia = 27.2-22.9 Mya *versus* 24.5 Mya and 22.4 Mya. By contrast, the ages inferred for the families are generally much younger in Eizirik et al. (2010) (**Figure 4**), a result potentially explained by a poor species sampling (two species of Ursidae, three species of Canidae, etc.) and the low phylogenetic signal of nuclear protein-coding genes for the most recent nodes.

The ages estimated for basal divergences in the Feliformia are much younger than in Eizirik et al. (2010) and Nyakatura & Bininda-Emonds (2012): Feliformia = 34.4-31.1 Mya *versus* 44.5 and 53.2 Mya; Feloidea = 27.4-24.7 Mya *versus* 33.3 and 52.9 Mya; and Viverroidea = 28.2-25.8 Mya *versus* 37.4 and 31.8 Mya. These results may be explained by the use of different calibration points: Feloidea = 34-20 Mya *versus* >28.5 Mya in Eizirik et al. (2010); Felidae = 20-14 Mya *versus* >31.15 Mya in Nyakatura & Bininda-Emonds (2012); and Viverridae = 34-14 Mya versus >25.72 Mya in Nyakatura & Bininda-Emonds (2012). The minimum age constraints used by Nyakatura & Bininda-Emonds (2012) for Felidae and Viverridae seem however problematic because they were not extracted from the fossil record, but from the supertree of Bininda-Emonds et al. (2007). By contrast, the minimum boundary of 28.5 Mya used for Feloidea in Eizirik et al. (2010) was interpreted from early Oligocene fossils assumed to be the oldest felids according to McKenna and Bell (1997). In Meredith et al. (2011), who used a similar minimum boundary for Feloidea (> 28.3 Mya), the divergence times estimated for Feliformia were found to be highly variable between their eight molecular dating analyses, with mean ages between 35.81 and 43.07 Mya. Three early Oligocene fossils were used as calibration points in this study: *Proailurus* and *Stenogale*, which were assumed to be stem Felidae; and *Palaeoprionodon*, which was supposed to be a stem Prionodontidae. However, the phylogenetic positions of these fossils remain uncertain. Indeed, the three fossil genera formed the sister-group of Feloidea + Viverroidea + *Herpestides antiquus*† in the phylogenetic analyses of Solé et al. (2014; 2016). In the classification of Hunt (2001), *Palaeoprionodon* was included in the subfamily Prionodontinae with *Prionodon* and other extant viverrid genera, such as *Genetta* and *Pioana*. Although *Prionodon* was treated as a member of the family Viverridae by Hunt (2001) and previous authors, it was then found to be the sister-group of the Felidae by Gaubert & Veron (2003) which was confirmed by more recent studies (e.g. Eizirik et al., 2010; Nyakatura & Bininda-Emonds, 2012; Meredith et al., 2011; **Figure 2**). In Nyakatura & Bininda-Emonds (2012), *Paleoprionodon* is used as a calibration point for the Viverridae (based on Hunt & Tedford, 1993, according to Bininda-Emonds et al., 2007), although it is suggested to be either close to *Prionodon* or a stem Feliformia (see e.g. Hunt, 2001; Veron, 2010; Solé et al., 2014).

Our dating estimates suggests that the basal split between *Nandinia* and other genera of Feliformia occurred at the Eocene/Oligocene transition (34 Mya), when a brutal and global cooling of 5°C resulted in the extinction of many taxa and the appearance of several modern mammal lineages (Russell & Tobien, 1986). Accordingly, *Nandinia* may be the descendant of forest-adapted ancestors, whereas the other lineage of Feliformia may have evolved in response to this important climatic change by adapting to more open vegetation.

## Conclusion

Based on a large taxonomic sampling (220 taxa represented by 2,442 mitogenomes) and 22 fossil calibration points (having both minimum and maximum age boundaries and a position based on recent fossil revisions and phylogenies), our study proposes a new time scaled phylogeny for Carnivora and provides further insights into the evolutionary history of this order. The age estimates for the Carnivora and the two suborders Caniformia and Feliformia fit well with global changes corresponding to the appearance of other mammal lineages. Moreover, our phylogenetic results, although largely similar to other recent phylogenies, suggest several taxonomic changes that would need to be confirmed using nuclear data.

## Acknowledgments

We thank our colleagues and their institutions for access to samples and for their help during this study: N. Chai and the staff of the Ménagerie du Jardin des Plantes, the staff of the Teuk Chhou zoo, H.E. Oknha Nhim Vanda, F. Catzeflis, R. Cornette, C. Denys, G. Dobigny, A. Delapré and the Mammal collection of the Muséum national d’Histoire naturelle, A. Délicat, J. Fuchs, P. Gaubert, C. Hatten, Y. Varelides, the Phongsaly Provincial Agriculture and Forestry Office and the EU Phongsaly Forest Conservation and Rural Development Project, S. Heard, A. Kitchener from the National Museums Scotland, J.-P. Hugot, S. Laidebeure and the staff of the Parc zoologique de Paris, E. Leroy, Van Cung Nguyen, P. Nicolau-Guillaumet, F. Njiokou, I. Parker M.-L. Patou, le Parc Zoologique de Montpellier, T. Petit and the staff of the zoo de La Palmyre, X. Pourrut, Peter Taylor and the Durban Natural Science Museum, the staff of Southport Zoo, Do Tuoc, B. Patterson, W. Stanley, and J. Phelps, from the Field Museum of Natural History, V. Volobouev, and L. Woolaver. This work was supported by the MNHN, CNRS, ‘PPF Biodiversité actuelle et fossile’, ‘Consortium national de recherche en génomique’, and the ‘Action Transversale du Muséum’ (ATM 2017: project RaPyD).

## Supporting Information

S1 Appendix. Origin of the sequences used in this study

S2 Appendix. Primers

S3 Appendix. Base composition

S4 Appendix. Distances

S5 Appendix. Bayesian tree reconstructed using the *mtDNA-Tv* dataset (220 taxa and 14,892 bp) and JC69+I+G model

S6 Appendix. Bootstrap 50% majority-rule consensus tree reconstructed from the MRP matrix of the SuperTRI analysis

S7 Appendix. Cytochrome b phylogeny of the genus *Melogale*

